# Metabolic composition of anode community predicts electrical power in microbial fuel cells

**DOI:** 10.1101/002337

**Authors:** André Grüning, Nelli J. Beecroft, Claudio Avignone-Rossa

**Affiliations:** *Corresponding Author*, Department of Computing, University of Surrey, Guildford GU7 2XH, United Kingdom.; Department of Microbial and Cellular Sciences, University of Surrey, Guildford GU7 2XH

## Abstract

Microbial Fuel Cells (MFCs) are a promising technology for organic waste treatment and sustainable bioelectricity production. Inoculated with natural communities, they present a complex microbial ecosystem with syntrophic interactions between microbes with different metabolic capabilities. From this point of view, they are similar to anaerobic digesters, however with methanogenesis replaced by anaerobic respiration with the anode as terminal electron acceptor. Bio-electrochemically they are similar to classical fuel cells where however the electrogenic redox reaction is part of the microbial metabolism rather than mediated by an inorganic catalyst.

In this paper, we analyse how electric power production in MFCs depends on the composition of the anodic biofilm in terms of metabolic capabilities of identified sets of species. MFCs were started with a natural inoculum and continuously fed with sucrose, a fermentable carbohydrate. The composition of the community, power and other environmental data were sampled over a period of a few weeks during the maturation of the anodic biofilm, and the community composition was determined down to the species level including relevant metabolic capabilities.

Our results support the hypothesis that an MFCs with natural inoculum and fermentable feedstock is essentially a two stage system with fermentation followed by anode-respiration. Our results also show that under identical starting and operating conditions, MFCs with comparable power output can develop different anodic communities with no particular species dominant across all replicas. It is only important for good power production that all cells contain a sufficient fraction of low-potential anaerobic respirators, that is respirators that can use terminal electron acceptors with a low redox potential. We conclude with a number of hypotheses and recommendations for the operation of MFCs to ensure good electric yield.

## 1 Introduction

In this paper we demonstrate a correlation between the power output of Microbial Fuel Cells (MFCs) and the composition of their anodic microbial communities with respect to metabolic function. This is important in order to understand the microbial functioning of MFCs and develop strategies to improve their reliability and yield. We focus on the analysis of the *metabolic* composition of the MFC community and the relation of community structure with power. This paper complements the analysis of MFC experiments in Beecroft et al. (2012) and Stratford et al. (2014) and, for the first time, is able to relate metabolic roles of various identified species with the power of a MFC.

MFCs are a promising technology for organic waste treatment and sustainable bioelectricity production (Logan and Regan, 2006, Lovley, 2006, Rabaey and Verstraete, 2005, Rittmann, 2006). Functionally, MFCs are bio-electrochemical reactors where microbes convert the chemical energy contained in organic substrates into electric energy by transferring electrons from the substrate directly or indirectly to the anode in the cell (Rabaey and Verstraete, 2005, Logan et al., 2006, Logan, 2009). Such fuel cells can be operated in batch-mode or in continuous mode (as here). Diverse MFCs designs and set-ups have been described with different anode and cathode arrangements (which we will not be concerned with in this article), and they can be fed with varying substrates, such as carbohydrate, acetate and other volatile fatty acids (VFAs), H_2_ or more complex and mixed feedstocks, for example wastewaters, or even more recalcitrant biomaterials such as lignocellulose and chitin (Huang et al., 2008, Ren et al., 2007, Rezai et al., 2009). They can be inoculated with a monoculture, or, as here, with a natural inoculum containing an unknown mix of species (Logan, 2009, Logan and Regan, 2006, Nevin et al., 2008). Different feedstocks and operating conditions will typically lead to the development of different microbial consortia (Kiely et al., 2011, Kouzuma et al., 2013), and even with identical starting and operating conditions consortia may diverge towards different phylogenetic compositions (Beecroft et al., 2012, Chae et al., 2009, Lee et al., 2008, Phung et al., 2004).

While genetic fingerprinting of the developing or mature consortia has been done occasionally (Velasquez-Orta et al., 2011, Katuri et al., 2011) to show that the consortium changes under different operating conditions, Beecroft et al. (2012), Kim et al. (2011) were among the first to fully analyse the composition of the anodic consortium during maturation in terms of its operational taxonomic units (OTUs). Experiments with and analysis of the microbial ecology of MFCs are – in addition to “technical” improvements to electrodes and membranes – important in order to increase their yield and stability by understanding the collective metabolism of a cell.

An MFC can be compared to a classical H_2_/O_2_ fuel cell where the electrogenic redox reaction is mediated by microbial species instead of by an inorganic catalyst. Like these, they consist of two compartments, the anode and the cathode, separated by a semipermeable membrane. Bacterial communities (or single species) attach to the anode electrode as biofilms. These constitute the biocatalyst for electricity generation. The electrons are transferred to the anode and travel around an external circuit (“load”) to the cathode while H^+^ cross the semipermeable membrane into the cathode chamber where they – in an air-breathing cathode like here – react with dissolved O_2_ and incoming electrons to form water.

An MFC can also be compared to an anaerobic digester where complex substrates are broken down to VFAs and other low molecular weight compounds along with H_2_ (acidogenesis) (Batstone et al., 2002). However the subsequent conversion of generated VFAs and H_2_ through methanogenesis is (mostly) replaced with anaerobic respiration where the anode is used as the terminal electron acceptor (Freguia et al., 2008, Lower et al., 2001). The anode thus serves two functions: it is the supporting structure of the biofilm and essential as terminal electron acceptor for respiration.

For complex substrates with natural inocula the community metabolism appears to be syntrophic between different species. For example the collective metabolism in carbohydrate-fed MFCs is essentially two-stage if inoculated with a multi-species mix of microbes (Freguia et al., 2008, Logan and Regan, 2006). In particular Freguia et al. (2008) found fermentation products such as acetate and H_2_ which were not present in the substrate, and they could also demonstrate that direct oxidation of the substrate did not happen to any significant extent. Hence, they concluded that the stage of fermentation is followed by a stage of anaerobic respiration using the anode as electron acceptor. In particular we understand the breakdown of substrate as follows: 1. Fermentation comprises all (anaerobic) processes breaking down the substrate to the generation of VFAs and H_2_. 2. Anode respiration is the anaerobic oxidation of VFAs and H_2_ with the anode as terminal electron acceptor (possibly with methanogenesis as a competing electron sink) (Lovley, 2008).

Metabolic stages might be complex: In a microbial consortium, fermentation might proceed step-wise with different microbes responsible for different metabolic steps, and respiration might involve several competing pathways such as respirative oxidation of VFAs or H_2_. Also, terminal electron transfer to the anode might proceed in a number of different ways (Rabaey and Verstraete, 2005, Bennetto, 1987, Reguera et al., 2005, Schröder, 2007, Gorby et al., 2006, Nevin et al., 2009, Kato et al., 2010), for example by: 1. direct electron flow from the respiratory enzymes in the microbial cell membrane in contact with the electrode, 2. extracellular organic or inorganic redox mediators that shuttle electrons between microbial cell and anode, or even 3. nano-wires, modified pili-like structures able to establish long range connections between microbial cells and terminal electron acceptors (such as metal oxides or graphite surfaces).

While Freguia et al. (2008) analysed very carefully the net metabolic reactions likely to play a role in electrogenesis in MFCs, they did not analyse how these might be distributed across different species. On the other hand, Beecroft et al. (2012), Ishii et al. (2012) found that the abundances of different microbes changed over time, while community function (that is power production) remained at comparable levels, and they did not find a consistent pattern of variation of individual species with cell power: It appears there is no single individual species facilitating good power generation in such MFCs.

In this paper we aim to clarify the functional metabolic structure of MFC communities. In our analysis, we assign the different OTUs found in the MFCs their most likely metabolic role based on the way they predominantly generate chemical energy in form of adenosin triphosphate (ATP) according to their known metabolism from literature, or that of closely related species. Our objective is to identify those metabolic capabilities a microbe needs to present in order to support good power production in a MFC. Our results show that respirative species able to grow against low anode potentials present the largest contribution to the power output of these MFCs.

## 2 Results

As reported in Beecroft et al. (2012), four replicate MFCs labelled *A, B, D* (6 data points each) and *C* (4 data points)^1^ were inoculated with anaerobic digester sludge. They were continuously fed with sucrose-containing media after the initial two week enrichment period for up to 91 days. Biomasses and identities of species as well as MFC power output were determined as described in Section 5.

### 2.1 Metabolic classification

We classified the metabolic function of identified OTUs into the following metabolic classes according to terminal electron acceptors:

**(f)** fermenters which use intracellular metabolites as electron acceptors.
**(r)** (anaerobic) respirators which typically fully oxidise fermentation products (VFAs and H_2_) with the help of an Electron Transport Chain (ETC). For later purposes we split (r) into two subclasses (Lengeler et al., 1999, Ch 12), see Table 1: **(n)** high-potential (anaerobic) respirators using electron acceptors such as O_2_ (in microaerophilics) or 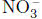, that is, acceptors with a relatively high redox potential. And **(s)** low-potential anaerobic respirators using electron acceptors with a comparatively low redox potential, such as 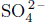.
**(o)** others, which can also be divided into two subgroups: **(a)** aerobic respirators which use O_2_, and **(u)** unknown.

**Table 1:**
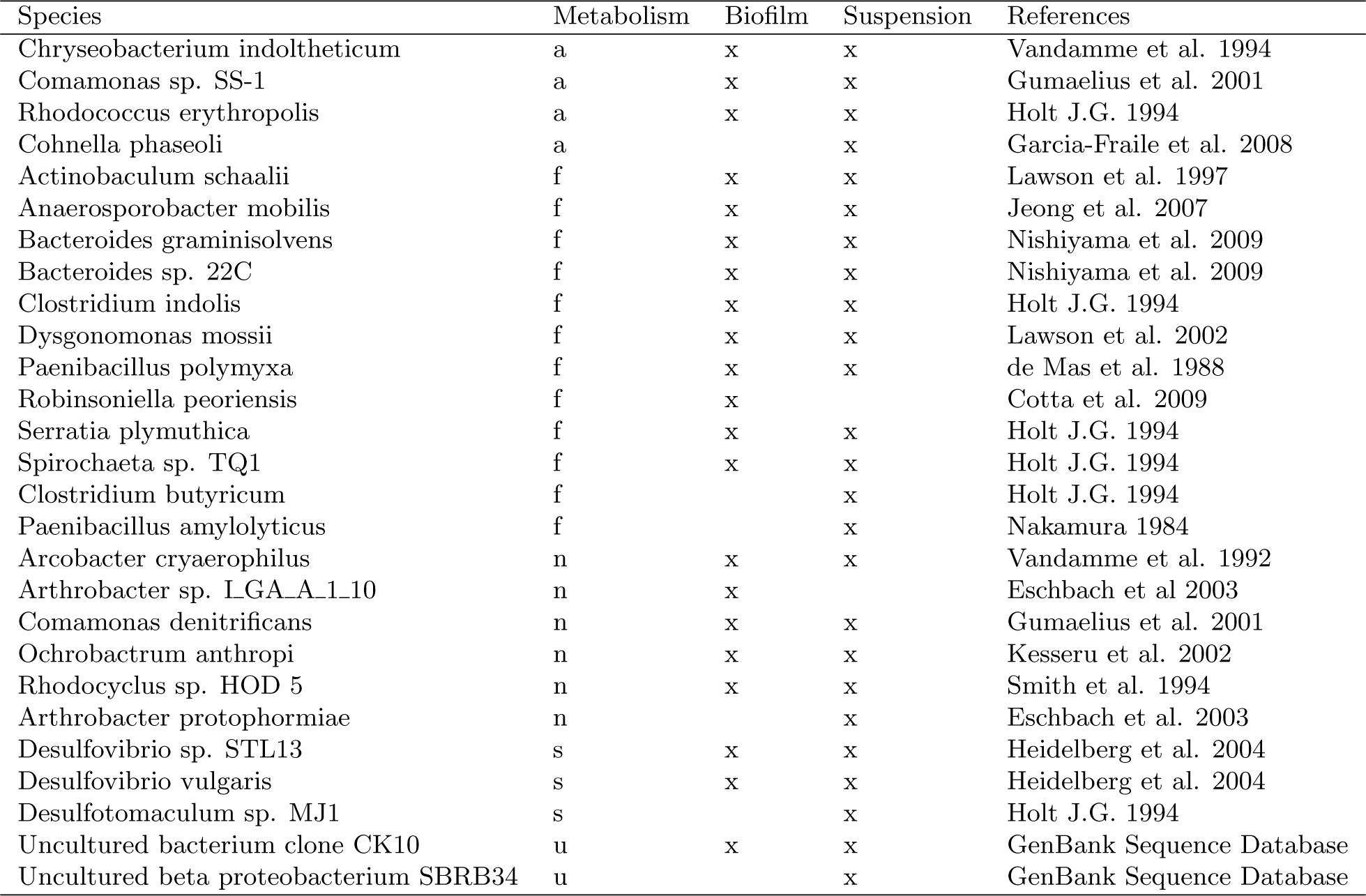
**Bacteria found in the biofilm and suspension of MFCs.** The table shows a classification of their most likely metabolic function: (a) aerobic respiration, (f) fermentation, (n) high-potential anaerobic respirators, (s) low-potential anaerobic respirators and (u) unknown, see main text for details.

The classification in Table 1 is based on the literature cited there; see Section 2.4 for how we dealt with cases where classification was ambiguous.

We concentrated initially on metabolic functions (f), (r), and (o) because an MFC with carbohydrate as substrate is essentially a two-stage system of fermentation (f) followed by anode-respiration (r) (Freguia et al., 2008). We assume that in principle each anaerobic respirator (r) that grows in the MFCs is a potential, but not necessarily good, electrogen. Finally we assume that aerobes (a) and species with yet unknown (u) metabolic pathways have only the effect of diverting electrons from the anode.

We added up the biomasses of all microbes with the same metabolic class (f),(r) and (o) and used these sums as our primary variables to describe the community. These measures of biomass per metabolic function (see Supplementary Table 7) are our primary variables in the subsequent analysis along with the age of a community (in days since start of continuous mode operation) and cell ids *A, B, C, D* as control variables external to the community).

Note that the MFCs are heavily anode-restricted (see Supplementary Information A). The relative anode restriction means that we are testing a system that is essentially not substrate-limited. We can hence directly see the consequences of limited anode-availability in a nutrient saturated community, that is we study only the effects of competition for the anode as electron acceptor, and not competition for the substrate.

### 2.2 Correlations

We determined the correlations of MFC power output with various system variables (Table 2) and we also confirmed that power does not vary significantly with MFC cell id (ANOVA, *p* = 0.288). Power increases significantly with age of the community. Hence a form of maturation or adaptation of the microbial community in the MFCs takes place through the run-time of the experiments. Also the biomass of respirators in the biofilm shows a strongly significant correlation with MFC power, in contrast with all other tested variables. It is not unexpected that the anaerobe respirators in the biofilm have a higher correlation with power than in the suspension: only those in the biofilm can donate electrons to the anode easily, and hence contribute to power. The considered MFCs are nutrient-saturated and almost all carbohydrates fermented to VFA and H_2_ (see Supplementary Table 8), and hence they would also be saturated in these two fermentation products. Thus much more VFA (or H_2_) substrate is available for the respirators than needed to achieve the theoretically maximal current through the load (see Supplementary Information A), and it is expected that power does not depend much on the fermenter biomass. Interestingly the correlation of the fermenters in the biofilm (albeit not significant) is greater than that of those in the suspension. A convenient interpretation of this is as a secondary effect: because the biomass of respirators goes up with power, these will also consume more VFA and H_2_ relieving negative feedback on the fermenters close to and in the biofilm in an otherwise VFA saturated suspension.

**Table 2:** **Correlations of power with other variables.** Correlations of power with age of the community in days and the biomasses of the different metabolic groups in the biofilm and suspension. Shown are Pearson correlation coefficients *r* along with the significance *p* of the correlation.

|  | r | p |
| --- | --- | --- |
| Day | 0.772 | 1.06e-04 |
| Respirators (film) | 0.806 | 3.15e-05 |
| Fermenters (film) | 0.225 | 3.54e-01 |
| Others (film) | 0.134 | 5.85e-01 |
| Respirators (susp) | 0.235 | 3.32e-01 |
| Fermenters (susp) | -0.142 | 5.63e-01 |
| Others (susp) | -0.173 | 4.78e-01 |

### 2.3 Linear models

From this point there are a number of natural paths to proceed in order to construct a linear model:

1. The Complete Model, which uses all available variables: the six biomass variables (namely the biomasses of fermenters (f), respirators (r) and others (o) in biofilm and suspension), the age (in days) of the community and cell id (A,B,C,D) to fit a linear model. This model (details Supplementary Table 9 and Supplementary Information B.1) has a correlation of *r*^2^ = 0.915 and hence models the power output very accurately with an Akaike Information Criterion (AIC) of 162.591, however at the cost of a high number of 11 parameters. Two further variables can be eliminated from this model, using step-wise reduction along the AIC (for details see Supplementary Information B.1).
2. The Community Model. By this we mean a linear model where we only included the variables that describe the composition of the community (that is all six biomass variables), but not those external to the community such as age and cell id. The model (see details in Supplementary Table 11 and Supplementary Information B.2 has a correlation of *r*^2^ = 0.762 and AIC 174.0798. We have here a noticeable decrease of *r*^2^ compared to the complete model (and subsequently also a marked increase of the AIC). This is not unexpected, and is an indication that the community variables on their own do not carry the full information about the adaptation towards power production. The only variable that is significant in the model (beside the intercept) is the biomass of respirators in the biofilm. This model can again be step-wise reduced along the AIC, and then has *r*^2^ = 0.747 and AIC of 169.1803 with only 4 parameters.
3. The Single Variable Model. Finally we can also use a model including only the biomass of the respirators in the biofilm as this is the variable that on its own has the strongest correlation with MFC power. Such a model has only two parameters left, and yields *r*^2^ = 0.649 and AIC 171.4307, see Table 3 and Figure 1. Compared to the initial complete model, the explained fraction of variance *r*^2^ has decreased, but this single variable model still captures about two thirds of the variance of the power data, which is remarkable.

**Figure 1:**
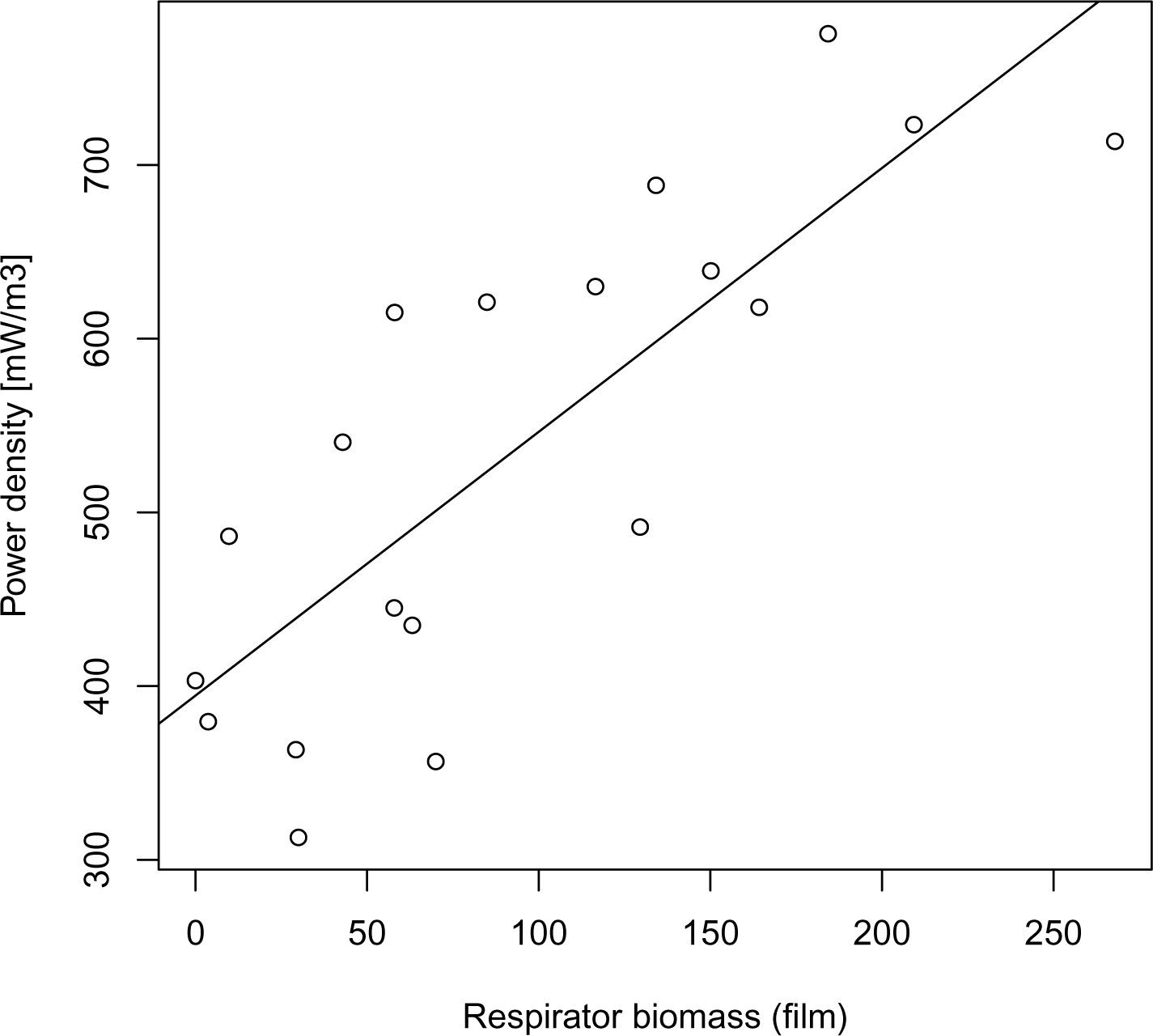
Linear regression of power as a function of respirator biomass [arbitrary units] in the biofilm.

**Table 3:** **Single variable model.** It contains the biomass of the respirators in the biofilm as its only variable.

|  | Estimate | Std. Error | t value | Pr(>\|t\|) |
| --- | --- | --- | --- | --- |
| (Intercept) | 395 | 32.5 | 12.1 | 8.5e-10 |
| ‘Respirators (film)’ | 1.52 | 0.271 | 5.61 | 3.15e-05 |

In the following we concentrate on this model with the single variable biomass of respirators in the biofilm. This is due to simplicity of presentation. In principle we could have chosen also the reduced community model or other linear models that include for example only the biofilm variables. Results for them [not reported] are similar to those reported below for the single variable model.

### 2.4 Metabolic classification

We classified OTUs in the MFCs according to their likely mode of energy metabolism into fermenters (f), anaerobic respirators (r) (and others) as described in Section 2.1. However information in the literature about the metabolism could be ambiguous. In those cases we dealt with the ambiguity as follows:

1. Where unambiguously clear from the literature, the microbe was assigned its unique role as fermenter (f), or respirator (r). For example *Comamonas denitrificans* cannot break down carbohydrates, but can oxidise short-chain organic acids with nitrate, so it is classified as (r) (Gumaelius et al., 2001).
2. Where a microbe is known to do both fermentation (f) and anaerobic respiration (r), it is typically classified as a fermenter (f) due to the competitive advantage of fermentation in high-density, high-nutrient environments, especially with limited access to external electron acceptors (Pfeiffer et al., 2001). For example *Serratia plymuthia* (here classed as (f)) is capable of 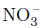 reduction, but also of fermentation (f) (Holt, 1994)
3. We further assumed that any anaerobic respirator (r) has the potential to be electrogenic due to the known wide flexibility of the ETC in prokaryotes, with a number of different electron entry and exit points (Logan et al., 2006). For example we classed *Rhodocyclus sp. HOD 5*, a very versatile purple non-sulfur bacterium (Smith et al., 1994), as (r) since it is capable of anaerobic respiration at least with 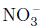 as terminal electron acceptor. However is capable also of anoxigenic photosynthesis which would be classed into the others (o).

We test below the effects of an alternative classification for the last two OTUs *S. plymuthia* and *Rhodocyclus sp. HOD 5*. These tests confirm our approach to class OTU into metabolic roles.

#### 2.4.1 *Serratia plymuthia*

*Serratia plymuthia* is capable of both fermentation (f) and 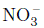 reduction (r) (Holt, 1994). According to our above argumentation about competition, *S. plymuthia* most likely acts as a fermenter. However we also retested the single variable model (Section 2.3) with *S. plymuthia* assigned role (r). In this case, the AIC sank to 175.3349 with a correlation *r*^2^ = 0.569. The result corroborates our criterion for classifying this OTU as a fermenter.

#### 2.4.2 *Rhodocyclus sp. HOD 5*

Similarly we looked at *Rhodocyclus sp. HOD 5*. This OTU is capable of 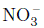 respiration but can also be anoxigenic photosynthetic and use a number of other pathways none of which seems to directly support electron transport to the anode more than (r) (Heidelberg et al., 2004). Assuming that this species did *not* contribute to anode respiration increased the AIC of the model to 173.5885 and decreased the correlation to *r*^2^ = 0.607. Hence it is more likely than not that *Rhodocyclus sp. HOD 5* acts as an anode respirator in our MFCs. The surprising importance of this species for current production in our MFCs is further corroborated by the lack of *Rhodocyclus sp. HOD 5* in an open circuit control experiment (Beecroft et al., 2012, Supplementary Material) and the results presented in Section 2.4.3 below.

#### 2.4.3 Systematically excluding potential anode respirators

We started from the hypothesis that all anaerobic respirators are potential electrogens. However we cannot be sure that all of them indeed act as anode respirators in our MFCs, or at least have a significant contribution in terms of rate of electron transfer to the anode towards current production in continuous mode operation with a relatively high load of 40kΩ. Hence for our model with a single variable left (Table 3), we systematically and iteratively took away individual respirators’ biomass from the respirator total biomass in the film – that is for the single variable model to effectively remove them from the model altogether. A removal was confirmed if it led to a higher correlation with power and reversed otherwise. Table 5 and Table 6 show the respirators whose biomass contributions were kept or excluded respectively towards correlation with the power.

The single variable model (details in Table 4) where biomass is only counted from the four OTUs in Table 5 has a correlation of *r*^2^ = 0.69 and AIC 169.1. This yields a very good fit with just one variable, and the correlation is higher than with the previous single variable model where *all* potential respirators were counted towards contributing to anode respiration.

**Table 4:** **Linear model of included species.** Linear model with only the *included* species counted towards respirator biomass in the biofilm, cp Table 5.

|  | Estimate | Std. Error | t value | Pr(>\|t\|) |
| --- | --- | --- | --- | --- |
| (Intercept) | 401 | 29.2 | 13.8 | 1.21e-10 |
| c(bbbb_respirators) | 2.45 | 0.398 | 6.15 | 1.08e-05 |

**Table 5:** **Included Species.** Respirators *included* after systematically swapping species out of class (r).

| Species | Metabolism |
| --- | --- |
| Arcobacter cryaerophilus | n |
| Desulfovibrio sp. STL13 | s |
| Desulfovibrio vulgaris | s |
| Rhodocyclus sp. HOD 5 | n |

## 3 Discussion

We discuss our results in order.

### 3.1 Correlations and linear models

We have tested a number of linear models containing subsets of the variables we have available. We lumped together biomasses of microbes with a comparable metabolic functions (fermenters (f), anaerobe respirators (r) and aerobic respirators and unknowns (o)). These models included a complete model that contains all variables and can explain about 90% of the variance of power (at the cost of 11 parameters), and a single variable model using only the respirator biomass in the biofilm as independent variable, using only 2 parameters, but still explaining about two-thirds of the variance of power between the four MFCs. For all models reported the respirators in the biofilm have a positive correlation with power in all models that do not include the community-external age as a variable.

### 3.2 Two metabolic stages

Our analysis supports the view of two-stage syntrophic metabolic processing (Freguia et al., 2008) with fermentation followed by anode-respiration. We assume that at any point in time each OTU participates in at most one process, either fermentation or respiration. This is suggested by cellular parsimony. For example the complete oxidation of sugars (an “in-house” combination of acidogenesis and respiration typically occurring during aerobic metabolism) does not happen prominently in MFCs, but is distributed across different OTUs. This is suggested by the typically higher rate of glucose uptake (but lower ATP yield) observed in fermentative metabolism as compared to aerobic respiration, and most probably to *anaerobic* respiration.

Accordingly, all OTUs whose joint biomass correlates strongly with power in Table 5 cannot use carbohydrate as a substrate (see literature in Table 1), but can oxidise only fermentation products anaerobically. Hence the electrogenic respirative stage relies on a fermentative stage that breaks down the carbohydrate to VFA and H_2_. We discuss the two stages in order.

#### 3.2.1 Fermentation stage

Fermentation works very reliably in our MFCs, and on average about 96% of carbohydrates are removed from inflow to outflow after day 14 during the continuous mode operation. Based on data (see Supplementary Table 8) from the experiments in Beecroft et al. (2012) there is no significant correlation after day 14 of the carbohydrate consumption with age (*p* = 0.232, correlation test), cell id (*p* = 0.984, ANOVA), or power (*p* = 0.423, correlation test). This may explain why power production does not depend strongly on the biomass in the fermentation layer with a correlation of *r* = 0.225*, p* = 0.354 vs a correlation of *r* = 0.806*, p* = 0.00315 × 10^−6^ for respirators, cp Table 2. This is also demonstrated by the lower significances of the fermentation variables within the linear models. Hence it seems that the *function* of the fermentation layer does not vary much over the time course of the experiment while Beecroft et al. (2012) did show that its *composition* does vary.

Fermentation is a faster process than respiration (Pfeiffer et al., 2001). If almost all sucrose is fermented to VFAs and H_2_ and – due to anode-restriction with a Coulombic efficiency of about 0.25% (see Supplementary Information A) – a low turn-over of fermentation products in the respiration layer, there must at all times be a relative excess of VFA and H_2_ substrate for the respirative layer. Thus a weak dependency of power on biomass in the fermentation stage is expected.

#### 3.2.2 Respiration stage

On the other hand, power should depend significantly on respirator biomass: Current is rate of electrons, and rate of electrons donated to the anode is rate of ETC rate times the number of individuals engaging in this process. Hence a strong dependence of current (and therefore power for a given load) on respirator biomass is expected. Although we are using linear models, it is unlikely that the dependency of power on biomass is linear for a wider range: doubling the respirator biomass will lead to less than double current, because with higher current the anode potential and consequentially the rate of ETC) sinks, because there is a higher voltage drop over the load. So we get less than double current through the load, but with a higher voltage drop, but these two factors are unlikely to compensate so as to leas to the same power.

For the following it will be useful to distinguish *high-* and *low-potential* respirators, ie those (anaerobic) respirators which can use terminal electron acceptors with a relatively high redox potential (such as O_2_ or 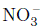) and those that can donate electrons to acceptors with a relatively low redox potential (such as 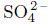).

##### High-Potential Respirators

The literature cited in Table 1 indicates that all excluded species in Table 6 come from genera or families where most members are aerobic or only facultatively anaerobic. And indeed also the excluded species themselves have an aerobic mode of respiration. Hence one could expect their anaerobic modes to be mainly reducing high-potential electron acceptors (such as O_2_, 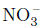 and other nitrogen compounds, see (Lengeler et al., 1999, Ch 10) for the redox potentials in the order of 0.3–1.3V in relevant reactions under physiological conditions). These OTUs are likely to be most competitive if a high-potential electron acceptor is available, or in an MFC if the anode potential is relatively high. Aerobic respiration is typically high-yield and low-rate (at least for O_2_ respiration, but likely also for other high-potential electron acceptors). That is, a significant amount of ATP are generated from each electron pair fed into the ETC, however at the cost of a low rate through a series of (slow) electron transport loops. Hence these organisms can gain considerable biomass while only contributing little to the current in the MFC.

**Table 6:**
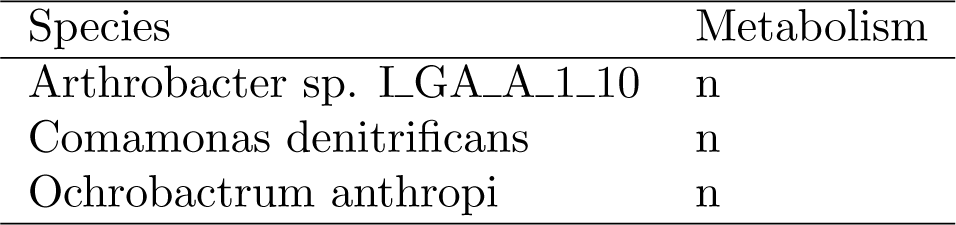
**Excluded Species.** Respirators *excluded* after systematically swapping species out of class (r).

##### Low-Potential Respirators

On the other hand, the *included* species in Table 5 come from genera presenting very diverse metabolic pathways, and none of them seems to have an outright aerobic mode, although they can be aero-tolerant or microaerophilic. The two *Desulfovibrio* species are known to be able to use 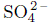 (and other sulfur compounds) as electron acceptors, while also purple (non-sulfur) bacteria such as *Rhodocyclus sp. HOD 5* have various pathways that involve inorganic sulfur compounds as electron acceptors (in dark reactions). Relevant reactions typically have redox potentials in the order of between −0.5V and −0.1V (Lengeler et al., 1999, Ch 10). Hence it is likely that all four included species can act as low-potential respirators with an ETC that can adapt to electron acceptors with low potential. This will be mainly at the cost of ATP produced per electron pair sent through the ETC (Lengeler et al., 1999, Ch 12). They most probably have a competitive advantage when only low-potential electron acceptors are available, or the anode potential is low, and they are likely to have a higher electron-transport rate than high-potential respirators if operated under low-potential conditions. It can therefore well be that a low-potential respirator is the driver of the current (due to high rate) still making only a comparatively minor fraction of biomass (due to low yield).

### 3.3 Bio-electrochemical conditions in the MFCs

The half-cell potential of the cathode *E_c_*, the current *I* through load *R* and the available redox potential *E_a_* at the anode are related as: *E_a_* = *E_c_ − U_L_*, where *U_L_*: = *RI* is the voltage drop across the load. This means that the anode acts as a terminal electron acceptor with a potential *E_a_* that gets lower at higher currents *I* through the load *R*. Let *E_s_* be the redox potential of the internal electron donor in the microbe (for example NADH or H_2_), then the *usable* potential *E_u_*: = *E_a_ − E_s_* = *E_c_ − E_s_ − RI* is proportional to the energy that the microbe can maximally retain from each electron passing through its ETC, and that is less than the potential difference *E_c_ − E_s_* between initial donor and terminal acceptor. The remainder *RI* has to be used to push the electrons into the anode and through the load. This means that the fraction of the energy contained in the substrate that the microbe can use for growth gets lower with increasing current through the load. Hence the ATP yield per e^−^ *must* sink with higher currents and lower anode potentials.

In continuous mode operation of the MFCs with a fixed 40kΩ load the measured voltage drop *U_L_* over the load is around or greater than 0.40 V (Beecroft et al., 2012). With an electrochemical half-potential of about *E_c_* = 0.82V in the air cathode, this yields an anode potential of around *E_a_* = 0.42V or less, *ignoring* any electrode over-potentials, hence the potential *E_a_* available to anode respirators is in the order of (or below) that of a typical 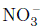 or 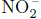 redox reaction. This might disadvantage high-potential respirators because the available potential is below the typical potential for high-potential respirators and give low-potential respirators a competitive advantage with a higher rate of electron transport, and hence a higher current through the load. As discussed above they might have a relatively high rate of electron donation while still producing little ATP and, consequently, biomass. See Logan (2009) for a similar argumentation.

#### 3.3.1 Succession of respirators

With a fixed load we should be able to observe that high-potential respirators are replaced by low-potential respirators as the biofilm matures (Rabaey and Verstraete, 2005): When the anode is colonised initially there is no current through the load, and therefore no voltage drop across the load from the cathode. This means that the first colonising species can utilise the full redox potential of the half-reaction in the cathode, ie *E_a_* = *E_c_* – in the case of an O_2_ cathode, this is the same potential as for aerobic respiration. Therefore the first colonising species can be high-potential respirators with high ATP-yield pathways and therefore high biomass yield, or low-potential respirators, which due to the high available potential show a higher rate of respiration and VFA and H_2_ consumption than normal.

However as the biomass of these initial colonists increases so does the rate of electrons donated to the anode. This means that the current through and the voltage drop over the load increase, leading to a lower potential of the anode. Therefore, in the course of time conditions become more favourable for low-potential respirators to dominate.

This argument is supported by experimental results because the joint biomass of the included group of bacteria of low-potential respirators in Table 5 correlates very well (*r* = 0.83*, p* = 0.00108 × 10^−6^) with power (and hence current) whereas the excluded group of high-potential respirators correlates less with power (*r* = 0.553*, p* = 0.014). The same picture emerges for the correlation of high- and low-potential respirator with the community age (*r* = 0.694*, p* = 0.0972 × 10^−6^ for low-potential respirators and *r* = 0.411*, p* = 0.081) for high-potential respirators).

### 3.4 H_2_ consumption and methanogenesis

Three out of the four included species are known to be able to consume H_2_ as substrate (*Rhodocyclus sp. HOD 5* and both *Desulfovibrio*). Molecular H_2_ is likely present in our MFCs as a fermentation product. With suitable ratios of concentrations of H_2_ and VFAs as potential substrates, the low-potential OTUs may consume H_2_ rather than VFA.

Methanogenesis (from acetate or H_2_) can be an attractive route to sink electrons besides anode respiration. However the pathways for methanogenesis are long, and their yield and growth rate of methanogens low (Lengeler et al., 1999, Ch 12). Methanogenesis gets more attractive for the consortium as a whole if the anode potential, and with it the available free energy for respiration, gets lower. However as long as there is a suitable electron acceptor (or by implication, the anode potential not too negative), low-potential respirators usually out-compete methanogens because their energy win is higher and they can grow at lower H_2_ concentrations and with a greater variety of carbon sources (Lengeler et al., 1999, Ch 12).

These considerations regarding competition for H_2_ (or acetate) between respirators and methanogens do not play a direct role in our substrate-saturated systems. They do however indicate that in anode-limited MFCs the available anode-potential must still be high enough so that anode respirators can still outcompete methanogens.

## 4 Conclusion

In this paper, we classified microbes found in MFCs according to their likely metabolic function based on the literature. We used biomasses per metabolic function as the primary model variables, and showed that power output varied strongly significantly and positively with the abundance of anaerobic respirators in the biofilm. Dependence on the fermenters was less pronounced.

Underlying the classification of microbes into classes fermenters (f) and anaerobic respirators (r) both contributing to MFC functioning, and others (o) which supposedly do not contribute or only negatively so, is the insight that an MFC with a fermentable substrate is essentially a two-stage syntrophic system (Freguia et al., 2008) with fermentation and respiration. In a septic system, there is no way to avoid fermentation that breaks down carbohydrates to VFAs and H_2_ due to faster metabolism of fermentation over respirators that hydrolyse complex substrates and subsequently oxidise them completely “in house”. Hence fermentation is done by one set of species, and complete oxidisation of resulting VFAs and H_2_ then “outsourced” to a group of anaerobic respirators. Ideally there is no electron acceptor except the anode and all electrons released in respirative oxidation are donated to the anode. Alternative electron sinks are oxygen contamination and methanogenesis, and finally biomass generation.

Within the respirative biomass, we identified a set four species whose joint biomass correlates most strongly with power. From known metabolic data about these species and their phylogenesis, it is likely that these four species are low-potential respirators. That is, they can grow competitively also when the anode potential is low. Other respirative species’ biomass varied less with power, and again from known metabolic data, it seems that these species are high-potential respirators that are most competitive when the anode potential is comparatively high. We conclude that for a mature anode biofilm with good electricity production, it is mainly the low-potential respirators that contribute to electricity generation. *Geobacter* has been reported widely to be important in current generation (Logan, 2009), however it seems to be just *one* low-potential respirator, and from our experiments other low-potential respirator exist that show a good performance in MFCs.

We conclude with a number of hypothesis resulting from our and others’ observations (Katuri et al., 2011, Rabaey and Verstraete, 2005, Logan, 2009) and considerations that do not yet seem to be reflected thoroughly in experimental work. If our observations and considerations regarding low- and high-potential respirators are valid, the following hypotheses pertaining to electric power production in MFCs emerge:

1. Electrogenesis is the norm, not the exception. That is, many more species than previously assumed can use the anode as electron acceptor (Kiely et al., 2011). These species are not necessarily good electrogens. However those that do best at a given anode potential will out-compete the others.
2. An MFC with complex substrate is a 2-stage system. That is, fermentable substrates are broken down into VFAs and H_2_ in a first step (”fermentation”). In a second step these are completely oxidised through respiration with the anode as terminal electron acceptor (”anode-respiration”). In a natural septic system with complex substrates, we cannot avoid having the fermentation stage as it would be more competitive to only do fermentation rather than hydrolysis/glycolysis and complete oxidation of its products “in-house”. Consequentially there will an unavoidable loss of chemical energy through fermentative organisms.
3. These two steps do not take place at the same time in a single organism, but are distributed across different species. Which role (fermenter or anode-respirator) a species capable of both takes on, depends on the prevalent environmental conditions in the MFC.
4. High external loads favour *low-potential respirators* that in nature grow against terminal electron acceptors with low redox potential (such as sulfate). High loads mean high voltage drops from the positive cathode to the anode, and hence lower anode potentials. Low-anode potentials (below 0.4V) mean that little energy can be generated in the ETC of the low-potential respirators, hence less chemical energy is retained in the microbes from the substrates, and more energy would be available to drive an external electric load. Anode potentials too low (below *−*0.2V to *−*0.4V, that is in the order of physiological redox potentials of H_2_ or NADH) will impede respiration so that methanogenesis becomes more competitive for H_2_ (and acetate) consumption than anode-respiration. This is a loss of available electrons. Optimal performance of a continuously operated MFC lies around that anode potential where methanogenesis is just not competitive against low-potential respirators. It is an open question whether keeping the anode potential constant (at a low voltage) or keeping the load fixed throughout colonisation yields a better electrogenic biofilm faster.
5. A MFC can only achieve a high Coulombic efficiency if the rate of substrate inflow and the ohmic load are in balance. The substrate is the electron donor, and hence its inflow rate determines the maximal current of the MFC in the case that all potentially available substrate electrons (for example 24 e^−^ for each sucrose) are ultimately donated to the anode. This electron donation can however only happen if the difference between the anode potential *U_A_* and that of the intracellular electron donor (typically NADH or H_2_) *U_S_* is positive, which in turn limits the maximal voltage drop *U_L_* across the load to the potential difference between the cathode reaction *U_K_* and the redox potential *U_S_* of the intracellular electron donor (in the order of a few V). This in turn determines the maximal load *R* = *U/I* for a given rate of substrate influx to achieve the maximal current for 100% Coulombic efficiency.
6. Measurement of polarisation curves to determine the load/current conditions of maximal power stem from classical electrochemistry. However their use in bioelectrochemical systems should be carefully considered: If an anode biofilm colonises the anode for a given fixed load or a given fixed anode potential, this biofilm will be acclimatised (for example regarding the fraction of low and high potential respirators) to the prevalent conditions (for example a high load). This means the community composition and metabolic activity is in balance with its environmental conditions at the end of maturation. If during the measurements of polarisation curves the external load is lowered (for some seconds or minutes) this measures only the short-term elasticity of the ETC rates of the current community (or possibly the rate of conversion of some intra-cellular storage polymer), and not the long-term point of maximal power of the community, as long-term (days to weeks) the community would have to acclimatise to the new load and change its composition accordingly. Hence measurement of the power of an MFC should happen under the same conditions as during maturation of the biofilm, as only this measures the long-term sustainable power of the cell. Hence our recommendations in this respect go beyond Menicucci et al. (2006), and are similar to those of Watson and Logan (2011) for batch operation: Set up identical MFCs, each with a different but fixed load throughout all acclimatisation, maturation and measurement of electric characteristics.

As always, this is the theory beset with practical detail to be tested in experiments.

## 5 Methods

### 5.1 MFC setup

The single-chamber MFCs contained carbon fibre veil anodes (32cm^2^) placed inside the anode chamber and connected to an electrical circuit with an insulated Ni/Cr wire. The air-breathing cathodes consisted of carbon cloth (9cm^2^) coated with Pt catalyst with polytetrafluoroethylene binder. A Nafion-115 proton-exchange membrane separated the anode chamber from the cathode.

The community composition and the peak power density were analysed between 14 and 91 days of operation. MFC voltage was monitored across a fixed external resistance of 40kΩ.

Originally, polarisation curves were recorded varying the external load in order to determine the peak power values. Here we use instead the power at the 40kΩ load fixed during maturation, because that load is in balance with the community structure (see Section 4). For the determination of the microbial community composition, the total deoxyribonucleic acid (DNA) was extracted from both a sample of the anode film (approximately 1cm^2^ out of a total surface of 32cm^2^) and 1 ml of the anode suspension (of a total of 7cm^3^ suspension volume). The partial bacterial (and archaeal) 16S rRNA genes were amplified by polymerase chain reaction (PCR) and analysed by Denaturing Gradient Gel Electrophoresis (DGGE). DGGE is a widely used molecular method for the semi-quantitative analysis of microbial community composition that gives reliable results for both environmental samples (Dong and Reddy, 2010, Ling et al., 2010, Wallis et al., 2010) and laboratory mixed cultures (Kongjan et al., 2010, Yang et al., 2012). It has also been used to demonstrate qualitative differences of community compositions (“community fingerprinting”) in MFCs but, unlike here, with out quantifying abundances, or identification of individual species (Katuri et al., 2011, Velasquez-Orta et al., 2011, Curtis et al., 2013).

Changes in the composition and structure of microbial communities have been monitored by using a combination of PCR-DGGE and DNA content analysis. Specifically, the relative proportion of species in the communities were inferred from the relative band intensities calculated by dividing the peak area of a band by the sum of peak areas of all bands in a lane (excluding chimeras). DNA was extracted from the DGGE bands and the species identified by sequence analysis. More details of the MFC setup and the methods used are described in Beecroft et al. (2012), Kim et al. (2011).

### 5.2 Data

The statistical analysis is based on the OTU abundances and MFC associated data of Beecroft et al. (2012).

Data points for cell/day A/41, B/56 and D/56 were excluded from the analysis because following a cathode change. Other data used are the power output of the MFCs at load 40kΩ along with the DNA concentration of the biofilm and the suspension. As the total DNA concentration times the ratio of sample size to total anode area or total suspension volume is a proxy of total biomasses, and the abundances from DGGE are only relative, we multiplied both to get a proxy of the biomass per OTU. This comes with two caveats: 1. The DNA concentration includes the DNA from the archaea. But as they are separated under different DGGE conditions, their relative abundance with respect to the bacteria is unknown. However methanogens are typically slowly growing due to their very low-yield metabolism, and are negatively affected by trace oxygen that may diffuse from the cathode. In similar experiments an archaea content of up to 20% was found (Chung and Okabe, 2009). 2. DNA content per cell varies for different bacteria. However DNA content has also been used successfully as a proxy for biomass in comparable settings (Kubota et al., 2009), so that DNA concentration times relative abundance is a somewhat distorted proxy of absolute species concentration. When we speak of “biomass” in this article, we understand it in the sense of the proxy as introduced above.

### 5.3 Statistical analysis

We test linear models that correlate the power output of our MFCs with a set of experimental variables describing the metabolic composition of the anodic community. As we have in the order of 20 data points and 21 OTUs in the biofilm and 25 OTUs in the suspension, a linear model that takes each individual OTU abundance as an independent variable is clearly over-specified. Furthermore Beecroft et al. (2012) did not find conclusive correlations of single OTUs with power. As a consequence we are aiming for simple models with only a few parameters, ideally less than four, but that takes account of the metabolic structure of the community.

Statistical analysis was done with R version 2.14.1 (2011-12-22) using the functions lm that can fit linear models with continuous and categorical variables such as cell id and step which stepwise eliminates variables to reduce the AIC.

## Acknowledgements

This work was supported by the Engineering and Physical Sciences Research Council grant number EP/I000992 (AG and CAR in part). NB and CAR were supported also by Research Councils UK Energy Programme as the Supergen 5 Biological Fuel Cells Consortium (managed by the Engineering and Physical Sciences Research Council: grants EP/D047943/1 and EP/H019480/1).

## Footnotes

1 This is the same numbering as in Stratford et al. (2014). Note however that cell *C* was not used in Beecroft et al. (2012), and cell *D* subsequently renumbered *C* in that paper.

## Supplementary Information

### A Coulombic Efficiency

For the experiments of Beecroft et al. (2012) we calculate the maximal current that would result from all available substrate electrons to be donated to the anode. According to Beecroft et al. (2012), the concentration of sucrose in the substrate solution is *c_m_* = 0.1 g*/*l, its molar mass is *m_n_* = 342.30 g*/*mol, and the volume rate with which the substrate solution is supplied is *r_v_* = 0.18 ml*/* min = 0.003 ml*/*s. Hence the molar rate of sucrose feed is *r_n_* = *r_v_ ∗ c_m_/m_n_* = 8.76*e −* 06 *×* 10^−12^mol*/*s. As each mole sucrose has 48 mol e ^−^ to donate, this is an inflow of e ^−^ at a rate *r_e_* = 48*r_n_* = 4.205*e −* 05 *×* 10^−9^mol*/*s. If all these electrons were donated to the anode, this would results in a current *I_max_* = *r_e_N_A_q_e_* = 4 mA where *N_A_* is the Avogadro constant and *q_e_* the elementary charge (which together are the Faraday constant *F* = *N_A_q_e_*).

If indeed all available substrate electrons passed through the anode and the load *R* = 40kΩ, this would results in a voltage drop of *U_L_* = *RI_max_* = 160V, and accordingly a difference of redox potentials between electron acceptor (O_2_ in the cathode) and electron donor (eg NADH or H_2_ to the ETC) in the same order of magnitude. This is electrochemically and physiologically impossible. The resistance of the load is too great to allow all electrons to pass through it because this would demand unrealistic redox potentials. Indeed with a load of 40kΩ, measured currents in the order of 5 − 15µA indicate a coulombic efficiency of about 0.25% (or 0.26% if we take into account that about 5% of sucrose is not consumed). This means, the MFCs used are anode-restricted with respect to the inflow of substrate, or alternatively are substrate-saturated with respect to the available electron sinks.

Assuming electrogenesis utilises an ETC with NADH or H_2_ as the electron donors (Kim et al., 2004), with physiological redox potentials in the order of *E_A_* = *−*0.32V for NADH/NAD or *E_A_* = *−*0.42V for H_2_/H (and in a similar order of magnitude for direct substrate oxidation), and the O_2_ cathode with *E_K_* = 0.88 V, the theoretical open-circuit voltage is *U* = *E_K_ − E_A_* ≈ 1.2V, (or for other electron donors and acceptors in the order of a few volts in any case). For a load *R* = 40kΩ this yields a maximal current of *I_max_* = *U/R* = 30µA if microbes did not retain any energy of the potential difference between the intracellular electron donor *E_K_* and the anode *E_A_* for growth or ATP generation. Measured maximal currents for 40kΩ are indeed in the same order of magnitude with 5 − 15µA, so that given the hard physical constraints of the load, cells achieve between 16%–50% of the physically maximally possible current. This is an efficiency in line with those measured in other experiments.

### B Details of linear models

#### B.1 Complete model

We used the six biomass variables (namely the biomasses of fermenters (f), respirators (r) and others (o)), the age (in days) of the community and cell id (A,B,C,D) to fit a linear model. This model includes all variables we have and is therefore called the *complete* model.

This model (details in Supplementary Table 9) has a correlation of *r*^2^ = 0.915 and hence models the power output very accurately with an AIC of 162.591, however at the cost of a high number of 11 parameters. We note that community age has a high significance which most likely reflects the increasing adaptation of the community to the environment in the cell. Cells do not differ significantly among themselves within this model.

Because of the high number of parameters, we were aiming for a model with fewer parameters which nevertheless has a good fit, and therefore removed variables iteratively if that reduced the AIC (using R’s step function). This led to model with AIC 158.68, *r*^2^ = 0.905 and 8 parameters. The 3 parameters removed were the biomasses of the fermenters in the suspension and those of the others both in the suspension and the biofilm, details in Supplementary Table 10.

#### B.2 Community Model

The complete model and its reduced version contain variables that represent external knowledge about the MFC such as the age of the community and cell id. We discuss here a model that is entirely based on the composition of the community including only the six biomass variables, see Supplementary Table 11.

The model has a correlation of *r*^2^ = 0.762 and AIC 174.08. We have here a noticeable decrease of *r*^2^ compared to the complete model (and subsequently also a marked increase of the AIC). This is not unexpected, and is an indication that the community variables on their own do not carry the full information about the adaptation towards power production.

We again reduce this model step-wise along the AIC (Supplementary Table 12). The reduced model has *r*^2^ = 0.747 and AIC of 169. We note that the significance of the respirators in the biofilm is strongest, followed the respirators in the suspension. These two along with the fermenters in the suspension are the only variables retained.

#### B.3 Model on only biofilm variables

Finally we test a model that includes only the biomasses of fermenters, respirators and others *in the biofilm* since in a substrate-saturated system the community in the biofilm should have the biggest effect on electron donation to the anode.

The results are summarised in Supplementary Table 13 with correlation *r*^2^ = 0.666 and AIC 174.468. In this model only the respirators are significant. A further step-wise reduction of variables along the AIC in Supplementary Table 14, leaves only a single variable, the biomass of the respirator. We have hence arrived at the single variable model, that in the main text we have selected based on the high correlation of the biomass of the biofilm respirators with the cell power.

**Table 7:**
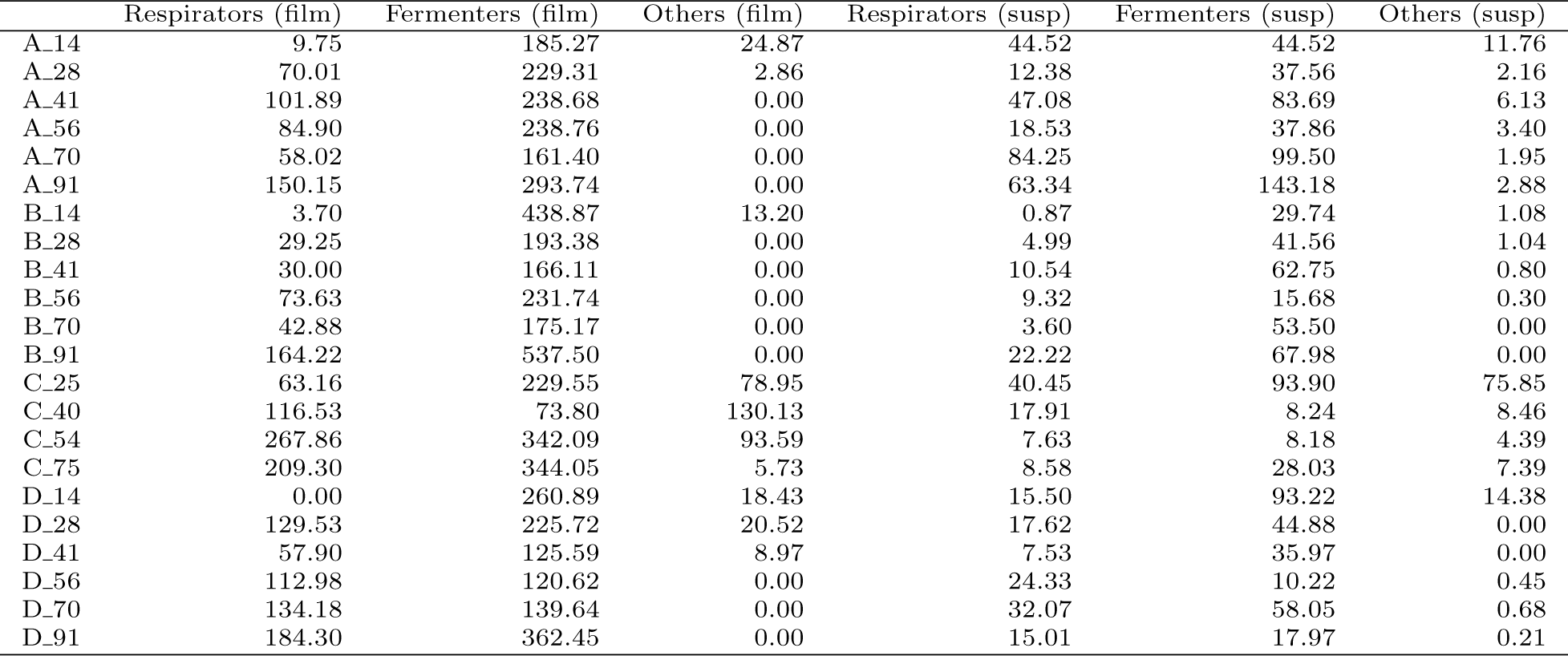
**Primary model variables.** Biomass per metabolic function and in arbitrary units (as described in the text). Data points are labelled A 14 etc., where A,B,C,D are the cell id, and the numbers denote the days since continuous mode operation started.

|  | Respirators (film) | Fermenters (film) | Others (film) | Respirators (susp) | Fermenters (susp) | Others (susp) |
| --- | --- | --- | --- | --- | --- | --- |
| A_14 | 9.75 | 185.27 | 24.87 | 44.52 | 44.52 | 11.76 |
| A_28 | 70.01 | 229.31 | 2.86 | 12.38 | 37.56 | 2.16 |
| A_41 | 101.89 | 238.68 | 0.00 | 47.08 | 83.69 | 6.13 |
| A_56 | 84.90 | 238.76 | 0.00 | 18.53 | 37.86 | 3.40 |
| A_70 | 58.02 | 161.40 | 0.00 | 84.25 | 99.50 | 1.95 |
| A_91 | 150.15 | 293.74 | 0.00 | 63.34 | 143.18 | 2.88 |
| B_14 | 3.70 | 438.87 | 13.20 | 0.87 | 29.74 | 1.08 |
| B_28 | 29.25 | 193.38 | 0.00 | 4.99 | 41.56 | 1.04 |
| B_41 | 30.00 | 166.11 | 0.00 | 10.54 | 62.75 | 0.80 |
| B_56 | 73.63 | 231.74 | 0.00 | 9.32 | 15.68 | 0.30 |
| B_70 | 42.88 | 175.17 | 0.00 | 3.60 | 53.50 | 0.00 |
| B_91 | 164.22 | 537.50 | 0.00 | 22.22 | 67.98 | 0.00 |
| C_25 | 63.16 | 229.55 | 78.95 | 40.45 | 93.90 | 75.85 |
| C_40 | 116.53 | 73.80 | 130.13 | 17.91 | 8.24 | 8.46 |
| C_54 | 267.86 | 342.09 | 93.59 | 7.63 | 8.18 | 4.39 |
| C_75 | 209.30 | 344.05 | 5.73 | 8.58 | 28.03 | 7.39 |
| D_14 | 0.00 | 260.89 | 18.43 | 15.50 | 93.22 | 14.38 |
| D_28 | 129.53 | 225.72 | 20.52 | 17.62 | 44.88 | 0.00 |
| D_41 | 57.90 | 125.59 | 8.97 | 7.53 | 35.97 | 0.00 |
| D_56 | 112.98 | 120.62 | 0.00 | 24.33 | 10.22 | 0.45 |
| D_70 | 134.18 | 139.64 | 0.00 | 32.07 | 58.05 | 0.68 |
| D_91 | 184.30 | 362.45 | 0.00 | 15.01 | 17.97 | 0.21 |

**Table 8:** Environmental data for the MFCs. *pH*, *P* power (density) in [mW*/*m^3^], concentration of DNA in film [*units*] and suspension, *U* voltage at fixed load 40kΩ, percentage of carbo-hydrate substrate consumed.

|  | Label | pH | P | DNA_film | DNA_susp | U | substrate |
| --- | --- | --- | --- | --- | --- | --- | --- |
| 1 | A_14 | 6.97 | 500.7 | 48.1 | 100.8 | 0.369 | 0 |
| 2 | A_28 | 6.95 | 408.1 | 66.1 | 52.1 | 0.316 | 95.85 |
| 3 | A_41 | 6.95 | 1578 | 74.5 | 136.9 | 0.491 | 99.05 |
| 4 | A_56 | 6.95 | 1048 | 70.8 | 59.8 | 0.417 | 98.28 |
| 5 | A_70 | 6.91 | 925 | 48 | 185.7 | 0.415 | 93.94 |
| 6 | A_91 | 6.91 | 1031 | 97.1 | 209.4 | 0.423 | 97.77 |
| 7 | B_14 | 6.99 | 393 | 99.7 | 31.7 | 0.326 | 0 |
| 8 | B_28 | 6.97 | 449 | 48.7 | 47.6 | 0.319 | 93.91 |
| 9 | B_41 | 6.96 | 356 | 42.9 | 74.1 | 0.296 | 94.94 |
| 10 | B_56 | 6.95 | 1054 | 66.8 | 25.3 | 0.402 | 93.85 |
| 11 | B_70 | 6.9 | 772 | 47.7 | 57.1 | 0.389 | 96.5 |
| 12 | B_91 | 6.94 | 1197 | 153.5 | 90.2 | 0.416 | 93.74 |
| 13 | C_25 | 6.88 | 433 | 81.3 | 210.2 | 0.349 | 37.54 |
| 14 | C_40 | 6.94 | 846 | 70.1 | 34.6 | 0.42 | 96.39 |
| 15 | C_54 | 6.91 | 1513 | 153.9 | 20.2 | 0.447 | 95.82 |
| 16 | C_75 | 6.94 | 1541 | 122.3 | 44 | 0.45 |  |
| 17 | D_14 | 6.97 | 474 | 61.1 | 123.1 | 0.336 | 37.42 |
| 18 | D_28 | 6.97 | 903 | 82.2 | 62.5 | 0.371 | 95.43 |
| 19 | D_41 | 6.96 | 756 | 42.1 | 43.5 | 0.353 | 98.27 |
| 20 | D_56 | 6.95 | 1645 | 51.1 | 35 | 0.468 | 97.63 |
| 21 | D_70 | 6.89 | 1591 | 59.9 | 90.8 | 0.439 | 95.65 |
| 22 | D_91 | 6.94 | 1792 | 119.6 | 33.2 | 0.466 | 94.04 |

**Table 9:**
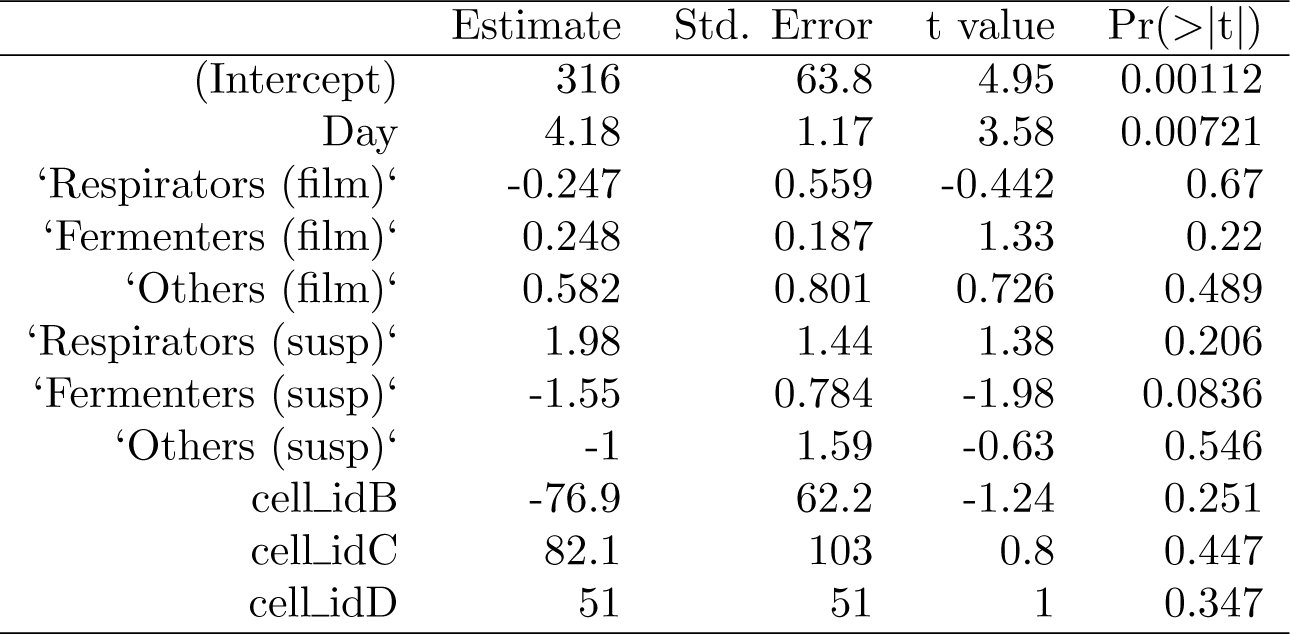
**Complete Model**. Estimates of parameters and their standard error, *t* statistic and the significance *p*.

**Table 10:**
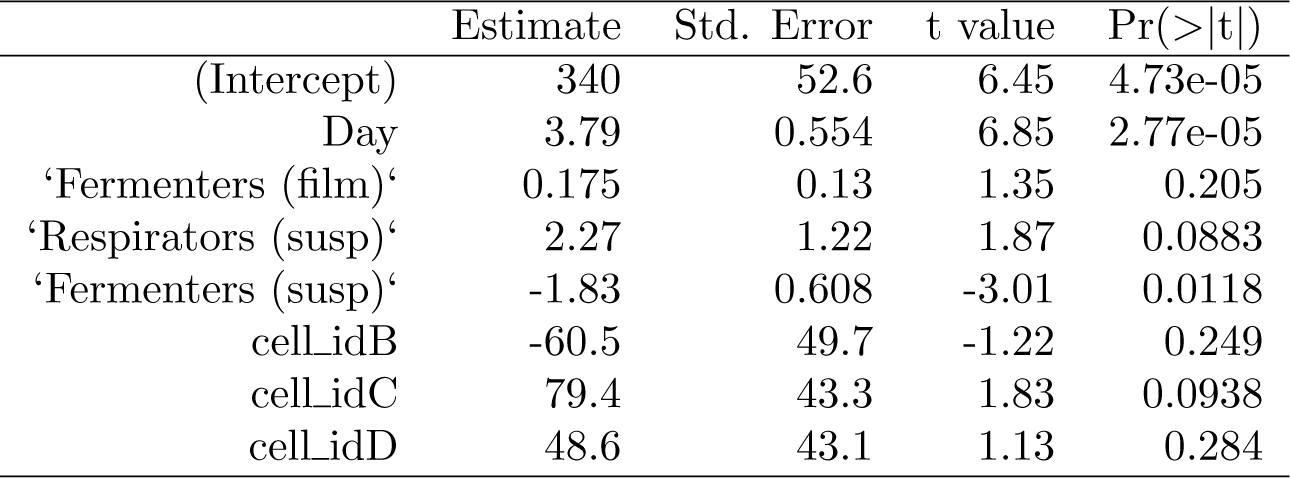
Reduced complete model

**Table 11:** **Community Model.** Biomass variables only

|  | Estimate | Std. Error | t value | Pr(>\|t\|) |
| --- | --- | --- | --- | --- |
| (Intercept) | 412 | 68 | 6.06 | 5.67e-05 |
| ‘Respirators (film)’ | 1.45 | 0.338 | 4.3 | 0.00104 |
| ‘Fermenters (film)’ | -0.0174 | 0.212 | -0.0818 | 0.936 |
| ‘Others (film)’ | -0.395 | 0.784 | -0.504 | 0.623 |
| ‘Respirators (susp)’ | 3.02 | 1.41 | 2.14 | 0.0531 |
| ‘Fermenters (susp)’ | -1.22 | 1 | -1.22 | 0.247 |
| ‘Others (susp)’ | -0.311 | 1.59 | -0.196 | 0.848 |

**Table 12:**
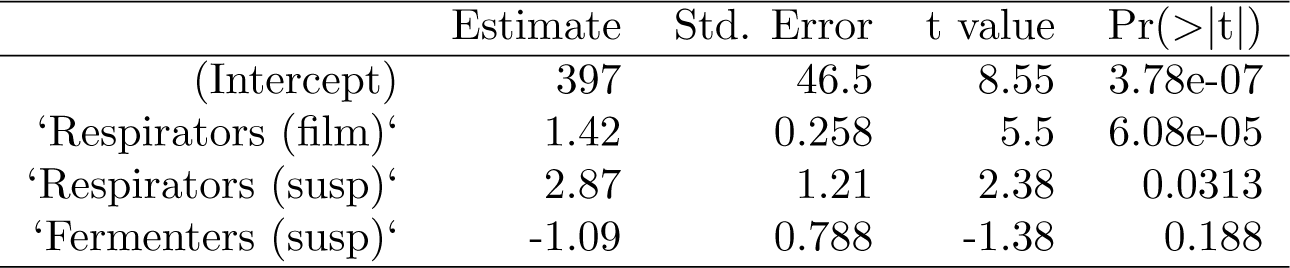
**Reduced Community Model.**

|  | Estimate | Std. Error | t value | Pr(>\|t\|) |
| --- | --- | --- | --- | --- |
| (Intercept) | 397 | 46.5 | 8.55 | 3.78e-07 |
| ‘Respirators (film)’ | 1.42 | 0.258 | 5.5 | 6.08e-05 |
| ‘Respirators (susp)’ | 2.87 | 1.21 | 2.38 | 0.0313 |
| ‘Fermenters (susp)’ | -1.09 | 0.788 | -1.38 | 0.188 |

**Table 13:** Linear model including only the three biomasses in biofilm.

|  | Estimate | Std. Error | t value | Pr(>\|t\|) |
| --- | --- | --- | --- | --- |
| (Intercept) | 430 | 54.6 | 7.87 | 1.05e-06 |
| bb_respirators | 1.66 | 0.325 | 5.12 | 0.000126 |
| bb_fermenters | -0.163 | 0.211 | -0.774 | 0.451 |
| bb_others | -0.402 | 0.616 | -0.653 | 0.524 |

**Table 14:** **Reduced Model of biofilm biomasses.** If the model with only the biofilm biomasses is reduced along the AIC we end up with the single variable model of the respirator biomass only.

|  | Estimate | Std. Error | t value | Pr(>\|t\|) |
| --- | --- | --- | --- | --- |
| (Intercept) | 395 | 32.5 | 12.1 | 8.5e-10 |
| bb_respirators | 1.52 | 0.271 | 5.61 | 3.15e-05 |

